# Pyramided resistance ensures grapevine (Vitis spp.) protection during high downy mildew (Plasmopara viticola) epidemic pressure

**DOI:** 10.64898/2026.02.11.705294

**Authors:** Guillaume Arnold, Tyrone Possamai, Emilce Prado, Emma Schlemmer, Sabine Wiedemann-Merdinoglu, Komlan Avia

**Affiliations:** INRAE, Université de Strasbourg, UMR SVQV, Colmar, France; Department of Crop Science, Research Institute of Organic Agriculture - FiBL, Frick, Switzerland

**Keywords:** Resistance breeding, Plant disease control, Downy mildew

## Abstract

Developing grapevine cultivars with genetic resistance to pathogens is a key strategy to reduce fungicide use and enhance sustainability. The French INRAE-ResDur program aims to pyramid several resistance loci against *Plasmopara viticola* (*Rpv*), the causal agent of downy mildew, while integrating factors against *Erysiphe necator* (*Ren/Run*) which causes powdery mildew. We evaluated in field the performance of grapevine genotypes carrying single or pyramided *Rpv* loci during the exceptionally severe downy mildew epidemic of 2024. Disease severity was quantified as the proportion of leaf foliage exhibiting symptoms. Susceptible controls averaged 66.6 % symptomatic leaves, *Rpv1/3.1* combination remained below 16.1 %. whereas the *Rpv1/Rpv3.1/Rpv10* pyramid showed only 4.9 % symptomatic leaves. The *s*ingle loci provided partial protection, but the effect varied with genetic background. Pyramiding improved resistance effectiveness and stability, indicating synergistic interactions among loci. These findings demonstrate that pyramiding *Rpv* loci is an effective strategy for durable downy mildew resistance and should be the preferred strategy in grapevine breeding programs and genetic resistance deployment strategies.

## Introduction

Cultivated grape varieties, primarily *Vitis vinifera*, are highly susceptible to major fungal diseases such as downy mildew, caused by *Plasmopara viticola*, and powdery mildew caused by *Erysiphe necator* (Bois et al., 2017; Gadoury et al., 2012; Gessler et al., 2011). These pathogens severely affect grape yield and quality, and under favorable conditions can cause complete yield losses (Fermaud et al., 2016; Gadoury et al., 2012; Gessler et al., 2011). The control of these diseases traditionally relies on repeated applications of chemical plant protection products (European Parliament, 2021; Innerebner et al., 2020). However, growing environmental and societal concerns have driven the viticulture sector to seek more sustainable alternatives (Fouillet et al., 2022; Pertot et al., 2017). Among the available solutions, breeding grape varieties with genetic resistance to fungal pathogens represents one of the most effective and sustainable approaches to reduce pesticide use (Merdinoglu et al., 2018; Mora et al., 2023; Trapp et al., 2025; Zhang et al., 2025). Over the past decades, numerous breeding programs, particularly in Europe, have intensified efforts to introgress resistance factors (loci) from wild *Vitis* species into cultivated *V. vinifera*. As a result, to date, 37 resistance loci to *P. viticola* (*Rpv*) and 20 loci to *E. necator* (*Run* or *Ren*) have been identified (Rockel, 2025). The effectiveness of single resistance loci varies widely, from partial to complete resistance (Possamai & Wiedemann-Merdinoglu, 2022), but several have already shown vulnerability to pathogen adaptation and resistance breakdown (Paineau et al., 2022; Pelissier et al., 2025; Peressotti et al., 2010). To improve the efficacy and durability of genetic resistance, multiple resistance loci can be combined within a single genotype, a breeding strategy known as ‘pyramiding’ (Mundt, 2018). Pyramiding increases the overall resistance level and decreases the likelihood that pathogen populations will simultaneously overcome all resistance factors, thereby enhancing the resistance durability (McDonald & Linde, 2002; Mundt, 2018).

In France, the INRAE-ResDur grapevine breeding program aims at developing new varieties carrying multiple resistance factors against *P. viticola* and *E. necator* (Avia et al., 2023; Merdinoglu et al., 2018; Mestre et al., 2013). The program mobilizes several key loci: *Rpv1* (Merdinoglu et al., 2003) and *Run1* (Pauquet et al., 2001), from *Vitis rotundifolia*; *Rpv3* (Bellin et al., 2009; Di Gaspero et al., 2012), *Ren3* (Welter et al., 2007) and *Ren9* (Zendler et al., 2020) from American *Vitis* spp. and *Rpv10* (Schwander et al., 2012) from the Asian species *Vitis amurensis*. All INRAE-ResDur populations carry *Run1*, conferring complete resistance to *E. necator* (Possamai & Wiedemann-Merdinoglu, 2022), as well as *Ren3* and *Ren9* which confer partial resistance. The ResDur generations differ by their *Rpv* combinations: ResDur1 combines *Rpv1* and *Rpv3*.*1*; ResDur2, *Rpv1, Rpv10* and ResDur3, *Rpv1, Rpv3*.*1* and *Rpv10*. Additional factors, such as *Rpv3*.*3* (Vezzulli et al., 2019), and the *Rgb1/Rgb3* loci conferring resistance to *Guignardia bidwellii* (Black Rot) (Bettinelli et al., 2023; Rex et al., 2014), have also been incorporated in recent generations. Most of the *Rpv* loci provide partial resistance to downy mildew (Possamai & Wiedemann-Merdinoglu, 2022), raising questions about the cumulative effectiveness of different pyramiding combinations, in vineyard conditions.

Since its launch in 2000, the INRAE-ResDur program has established several field selection plots at the INRAE center in Colmar, where genotypes carrying single and combined *Rpv* factors are maintained (Schneider et al., 2019) . The exceptionally wet and mild 2024 season created ideal conditions for *P. viticola* infection across France, leading to a historically severe downy mildew epidemic. This provided the opportunity to assess, under a high natural disease pressure, the in-field performance of grapevine genotypes carrying different combinations of single and pyramided *Rpv*. Our results demonstrate that pyramiding substantially enhanced resistance effectiveness and stability compared to single loci. These findings provide valuable insights into the design and deployment of durable resistance in grapevine breeding programs.

## Materials and methods

### Rpv genotyping

The presence of *Rpv* loci in the INRAE-ResDur grapevine populations was determined using simple sequence repeat (SSR) markers following the protocol of Blasi et al. (2011), with minor modifications. Genomic DNA was extracted from young leaf tissue using the DNeasy 96 Plants DNA kits (Qiagen, Hilden, Germany), according to the manufacturer’s instructions. SSR amplification was performed with 5’ fluorescently labeled primers (FAM, HEX or NED; Applied Biosystems-Thermo Fisher Scientific, Waltham, MA, USA). Following a 1:5 dilution in nuclease-free water, 1 µL of each PCR product was mixed with 11 µL of formamide containing the internal standard HD400-ROX (Applied Biosystems). Fragment separation was carried out via capillary electrophoresis on an ABI Prism 3100 Genetic Analyser (Applied Biosystems, Thermo Fisher Scientific, Waltham, MA, USA) equipped with 50 cm capillaries and POP-4 polymer. Electropherograms were analyzed using GeneScan™ 3.1 software (Applied Biosystems), and the SSR alleles were identified with Genemapper™ (Applied Biosystems). The presence of *Rpv1, Rpv3*, and *Rpv10* loci was confirmed by using at least two flanking SSR markers per locus (Table 1; Supplementary File S1): VMC4f3.1 and VMC8g9 for *Rpv1* (Barker et al., 2005); SC8-0096-022 (Genoscope, France), UDV737 (Di Gaspero et al., 2012) and VMC7f2 (Bellin et al., 2009) for *Rpv3*.*1 and Rpv3*.*3*; Gf09-44 and Gf09-47 for *Rpv10* (Schwander et al., 2012). Genotypes were considered positive for a given *Rpv* locus only when both alleles were consistent with the resistant parental genotype and non-recombinant for the two flanking markers. Individuals with incomplete or inconsistent genotyping data were excluded from the study.

**Table 1.** Details of the SSR markers utilized in the *Rpv* genotyping. Resistance loci to *P. viticola* (*Rpv*) segregating in the INRAE ResDur populations, genetic SSR markers utilized for the assisted selection, and alleles associated with the resistance. *Rpv3s* loci colocalize in the grapevine genome but are associate to different alleles. Complete parental plants allelic profile in Supplementary File S1.

| ***Rpv* factor** | **SSR marker** | **Resistance-linked allele** |
| --- | --- | --- |
| *Rpv1* | VMC4f3.1 | 189 or 191 |
| *Rpv1* | VMC8g9 | 155 |
| *Rpv3.1 / Rpv3.3\** | SC8-0096-022 | 220 / 222* |
| *Rpv3.1 / Rpv3.3\** | UDV737 | 279 / 271* |
| *Rpv3.1 / Rpv3.3\** | VMC7f2 | 207 / 197* |
| *Rpv10* | Gf09-44 | 230 |
| *Rpv10* | Gf09-47 | 296 |

### Plant material

A total of 286 grapevine were evaluated from four segregating populations of the INRAE-ResDur breeding program, selected for the presence of *Rpv1, Rpv3*, and *Rpv10* resistance loci (Supplementary File S2). Each population represented specific combinations of resistance loci, with 8-41 genotypes per combination. The population 50001 (ResDur1 generation) was originated from a cross between the *V. rotundifolia* backcross line ‘3082-1-42’ (carrying *Rpv1*) and the variety ‘Regent’ (carrying *Rpv3*.*1*). The population 42050 (ResDur2 generation) resulted from a cross between ‘3160-11-3’, a *V. rotundifolia* backcross carrying *Rpv1*, and ‘Bronner’ which carries *Rpv3*.*3* and *Rpv10*. The population 50035 (ResDur3 generation) was obtained by crossing ‘Artaban’ (ResDur1 generation carrying *Rpv1* and *Rpv3*.*1*) with ‘Divico’ (*Rpv3*.*3* and *Rpv10*). A fourth, mixed population, including 77 genotypes derived from various ResDur2 and ResDur3 crosses, integrating diverse *Rpv* combinations. All populations segregated for multiple *Rpv* loci and shared *Run1*-based resistance to *Erysiphe necator*, except *Rpv-* susceptible genotypes of 50001 population.

### Vineyard conditions and experimental design

All populations were cultivated at the INRAE experimental vineyard in Colmar, France, on deep loess soils (Supplementary File S3). Genotypes were randomly distributed within each plot, with four adjacent vines per genotype. Vines were trained to Rhine espalier system with double Guyot pruning, at the row spacing of 1.7 m and an intra-row spacing of 1.4 m, resulting in a planting density of 4,200 vines per hectare. Populations 50001 and 42050 were planted in 2004 and 2008, respectively, grafted onto Fercal (clone 242) roostock. Population 50035 was planted in 2017 onto SO4 (clone 742). The mixed population (77 genotypes) was planted in 2011, also grafted onto SO4 (clone 742). Until 2021, the experimental plots received no fungicide treatments. Beginning in 2022, minimal phytosanitary applications were introduced to control Black Rot, powdery mildew and *Pseudopezicula tracheiphila* (Rotbrenner). During the 2024 growing season, treatments were applied (3 June and 21 June). Meteorological data (precipitation and temperature) for 2024 were recorded at the Colmar weather station and integrated into the INRAE AgroClim Avignon network for environmental characterization (Delannoy et al., 2022).

### Assessment of downy mildew damages on leaves

Downy mildew severity was evaluated on 2 September 2024, following the epidemic phase. For each genotype, the proportion of symptomatic leaf area was visually estimated based on leaves damages induced by a direct effect of *Plasmopara viticola* infection such as defoliation, dessication of the leaf blade, sporulation but also by necrosis associated with grapevine defence response. One categorical score for each genotype (four vines) was assigned according to the estimated percentage leaf area with symptoms: 0%, 1%, 2.5%, 5%, and subsequently in 5% increments (Supplementary File S5). This scoring system captured the estimated extend of leaf damage or disease severity, under field epidemic conditions.

### Statistical analysis

All statistical analyses were conducted in R v4.3.1 (R Core Team., 2023). The Kruskal-Wallis test was used to detect significant differences in downy mildew severity among *Rpv* combinations, followed by Dunn’s post hoc test. P-values were adjusted using Benjamini-Hochberg false discovery rate (FDR) correction with a significance threshold of alpha = 0.05 (Ogle et al., 2023). This non-parametric approach due to heteroscedasticity and unequal sample sizes among genetic classes. Graphical data representation was performed using ggplot2 package (Wickham, 2016)

## Results

### Meteorological conditions and epidemic development

The 2024 growing season in Colmar was characterized by persistent rainfall and moderate temperatures, which created highly conducive conditions for *P. viticola* infections (Figure 1). The first infection event occurred on 2 May 2024, when cumulative rainfall exceeded 10 mm and the mean temperature was above 10°C, favorable thresholds for primary inoculum release. The first *P. viticola* sporulations were detected on 13 May 2024, followed by multiple overlapping secondary cycles extending though late summer. Consequently, the 2024 epidemic represented one of the most severe downy mildew outbreaks recorded in the Alsace region (Chambre d’agriculture d’Alsace et Fredon Grand Est, 2024).

**Figure 1.**
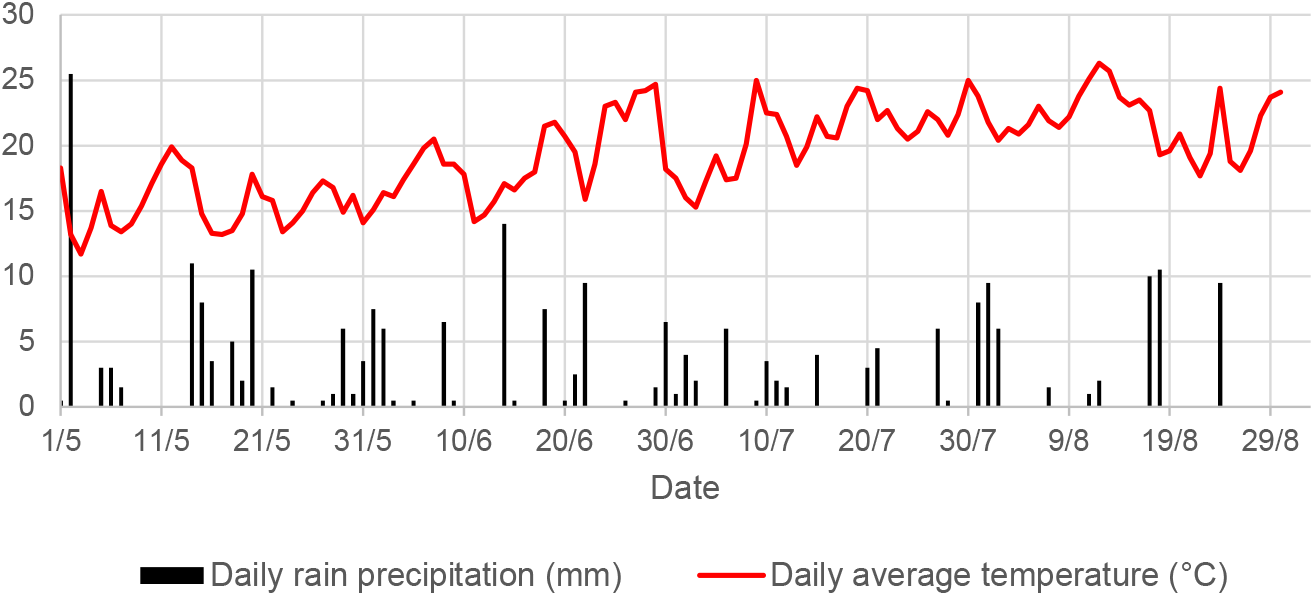
Weather conditions in 2024 at INRAE in Colmar. Average daily temperature (°C; red) and rainfall (mm; black), which favored the spread of the downy mildew epidemic at INRAE-UMR 1131 SVQV (Colmar, France), from the first occurrence of suitable conditions for primary infections (01/05) until the day on which disease symptoms were assessed in the field (02/09).

### Performance of Rpv combinations across populations

The field assessment revealed significant differences in downy mildew severity among the studied *Rpv* combinations (Kruskal–Wallis test, p < 0.05; Figure 2). In all populations, genotypes carrying pyramided resistance loci (more than 1 locus) exhibited substantially lower symptom severity than those carrying single *Rpv* factors or none (Figure 2).

**Figure 2.**
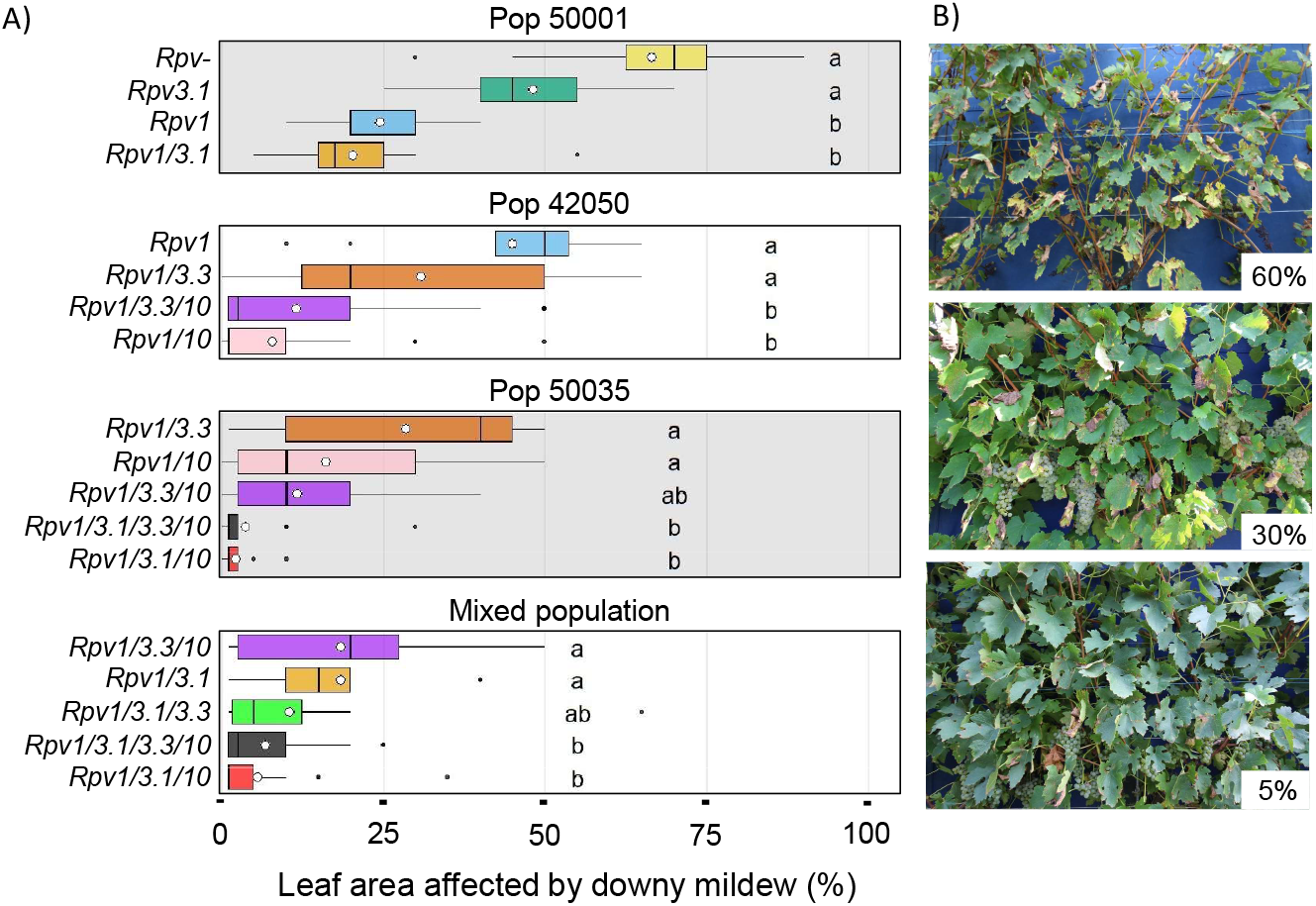
Downy mildew damages on the INRAE ResDur grapevine populations. A) Distribution (box plots) and mean (white dot) of leaf area affected by downy mildew (%) for the *Rpv* combinations (coloured) of the different study populations (50001, 42050, 50035 and mixed population). Between 8 and 41 genotypes per *Rpv* combination were assessed. Comparisons within each population were carried out using a Kruskal-Wallis test supplemented by Dunn’s post hoc tests (significant differences for p-values < 0.05). Data details in Supplementary File S4. B) Photos representing three levels of leaf area affected by downy mildew (%) according to different level of resistance: 60%, 30%, and 5%. Complete range of symptoms and further levels of infection are shown in Supplementary File S5.

#### Population 50001 (ResDur1)

This population segregated for *Rpv1* and *Rpv3*.*1*. Susceptible controls (*Rpv*-) showed, on average, 66.6% symptomatic leaf area, whereas *Rpv3*.*1* monogenic genotypes, showed moderate but non-significant reductions of symptoms to 48.2 %. Genotypes carrying *Rpv1* alone had significantly lower symptomatic leaf area (24.4 %), and the *Rpv1/3*.*1* pyramided combination showed an even greater reduction of symptoms (20.4 %), suggesting additive effects between loci.

#### Population 42050 (ResDur2)

In the *ResDur2* generation, the combination of *Rpv1, Rpv3*.*3*, and *Rpv10* was evaluated. Monogenic *Rpv1* genotypes were the most susceptible, averaging 45% symptomatic leaf area. The *Rpv1/3*.*3* combination yielded a modest but non-significant reduction of symptoms (30.9 %). In contrast, the addition of *Rpv10* significantly decreased the disease severity to 11.6 % in *Rpv1/3*.*3/10* and 7.9 % in *Rpv1/10* genotypes, confirming the lowest severity and therefore the highest protective effect of *Rpv10* under epidemic conditions.

In an unexpected way, for the *Rpv1* combination, the percentage of leaf damaged was notably higher compared to the one observed on *Rpv1*-carrying genotypes of population 50001.

#### Population 50035 (ResDur3)

This population combined *Rpv1, Rpv3*.*1, Rpv3*.*3*, and *Rpv10* in various configurations. The *Rpv1/3*.*3* combination remained moderately affected by downy mildew (28.5% symptomatic leaf area) similarly to the disease severity observed in the 42050 population, whereas *Rpv1/10* and *Rpv1/3*.*3/10* genotypes exhibited significantly reduced infection levels (16.2% and 11.8%, respectively). The most effective combinations, *Rpv1/3*.*1/10* and *Rpv1/3*.*1/3*.*3/10*, showed important reduction of symptoms, with only 2.3 % and 3.8 % of the leaf area damaged, respectively. These results confirm the additive or synergistic contribution of *Rpv3*.*1* and *Rpv10* to resistance effectiveness.

#### Mixed population (ResDur2 + ResDur3 crosses)

Genotypes from various ResDur2 and ResDur3 generations exhibited resistance patterns consistent with those observed in structured populations. The *Rpv1*/*3*.*1* and *Rpv1/3*.*3/10* combinations conferred intermediate protection (18-19 % symptomatic area), while *Rpv1/3*.*1/10* and *Rpv1/3*.*1/3*.*3/10* combinations provided strong reduction of symptoms, with 5-7 % mean leaf damage. The *Rpv1/3*.*1/3*.*3* combination, which was not evaluated in the biparental populations, exhibited a moderate-to-high level of resistance, with an average of 10.6% of leaves affected, representing a 7.8% reduction in disease severity compared to *Rpv1/3*.*1* genotypes. Across populations, *Rpv3*.*3* contributed little additional benefit when combined with *Rpv1* or *Rpv10*, while *Rpv3*.*1* and *Rpv10* consistently enhanced field performance.

### Overall resistance patterns and comparative analysis

Pooling data across all populations carrying mains *Rpv* loci utilized in the INRAE-ResDur breeding program (i.e., excluding *Rpv3*.*3*) revealed a clear, hierarchical gradient of resistance effectiveness in pyramiding by reducing extend of leaf damages (Figure 3; Supplementary File S6). Susceptible genotypes (*Rpv*–) averaged 66.6 % leaf damage, while *Rpv3*.*1* and *Rpv1* alone reduced infection to 48.2 % and 31.5 %, respectively. Pyramided combinations further reduced disease severity due to improved resistance: *Rpv1/3*.*1* averaged 16.1 %, *Rpv1/10* averaged 12.3 %, and *Rpv1/3*.*1/10* showed the highest field resistance, with 4.9 % symptomatic foliage. Statistical analysis confirmed that all pyramided combinations involving *Rpv10* were significantly more resistant than single-locus genotypes (p < 0.001). These results demonstrate that resistance effectiveness increases with the number of stacked loci, with *Rpv10* providing the strongest incremental contribution under epidemic pressure.

**Figure 3.**
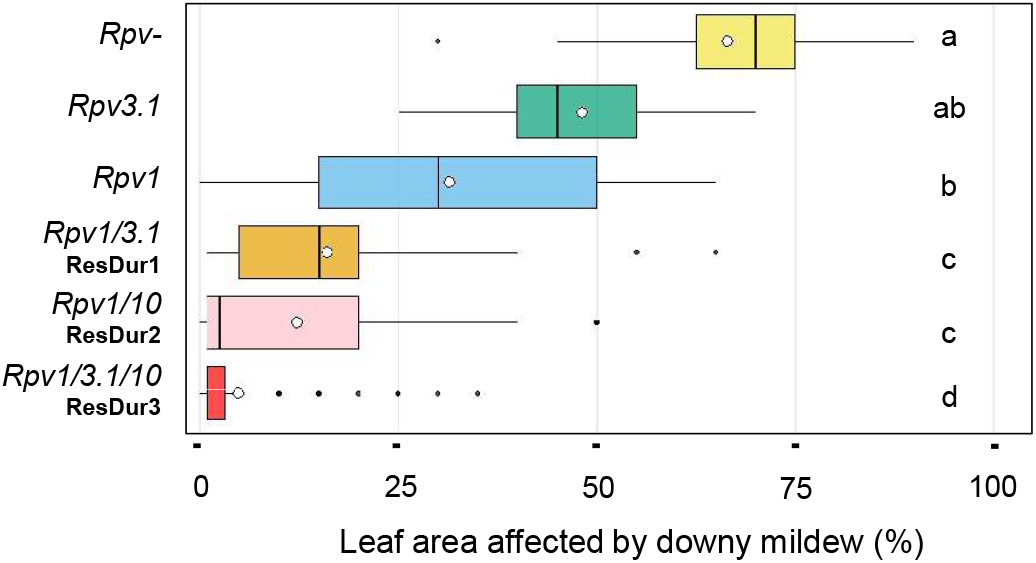
Downy mildew damages on the *Rpv* factors and combinations of the INRAE ResDur programme (excluding Rpv3.3) . Global distribution (box plots) and mean (white dot) of leaf area affected by downy mildew (%) for the mains *Rpv* combinations (coloured) of the INRAE ResDur breeding program. Data from the different populations were merged and comparisons were carried using a Kruskal-Wallis test supplemented by Dunn’s post hoc tests (significant differences for p-values < 0.05). Data details in Supplementary File S6.

## Discussions

### Field validation of Rpv pyramiding under epidemic conditions

The exceptionally rainy and mild 2024 season provided a unique opportunity to evaluate the effectiveness of *Rpv* loci combinations under natural epidemic pressure. Our findings confirm that resistance conferred by individual *Rpv* loci, particularly *Rpv1* and *Rpv3*.*1*, remains partial and variable, whereas pyramiding these loci significantly enhances protection against *P. viticola*. These results are consistent with previous controlled-environment studies (Calonnec et al., 2013; Possamai & Wiedemann-Merdinoglu, 2022; Possamai et al., 2026), and demonstrate that additive or synergistic interactions among *Rpv* loci can be also observed under complex and polycyclic field epidemics.

The superior performance of the *Rpv1/3*.*1/10* combination, showing less than 5% symptomatic leaf area even under extreme disease pressure, illustrates the success of the INRAE-ResDur breeding program’s pyramiding strategy. By contrast, single-locus genotypes displayed heterogeneous and unstable responses, underlining the limitations of monogenic resistance for durable field protection.

### Comparative contributions of Rpv loci and influence of genetic background and pathogen adaptation

The field data confirmed significant effects on grapevine resistance for *Rpv1, Rpv3*.*1*, and *Rpv10*, while *Rpv3*.*3* provided limited additional resistance. These findings align with prior laboratory assays performed at INRAE Colmar (Wiedemann-Merdinoglu et al., 2022; Chedid et al., 2023), where *Rpv3*.*3* showed weak or background-dependent effects. The absence of a detectable contribution in our field trials suggests that *Rpv3*.*3* alone does not confer substantial quantitative resistance in the tested backgrounds or against the prevailing *P. viticola* populations.

Notably, the resistance level conferred by *Rpv1* varied between populations 50001 and 42050, suggesting a host genetic background effect. Such variability may reflect the differential expression of downstream defense pathways or quantitative modifiers interacting with *Rpv* loci. The breakdown of mains *Rpv* utilized in breeding programs, including *Rpv1, Rpv3*.*1, Rpv10*, have been widely observed across Europe (Paineau et al., 2022).In our study, only the *Rpv3*.*1* resistance have been consistently eroded but the locus still conserved a significant effect when combined in pyramiding, suggesting still an incomplete breakdown of the locus.

Among the tested *Rpv* loci, *Rpv10* provided the most consistent and robust results in reducing downy mildew symptoms across populations when combined with *Rpv1* or *Rpv3*.*1*, evidencing the strategic value of this genetic resource in the INRAE-ResDur breeding program and future pyramids.

### Implications for resistance durability and breeding strategy

Notably, the pyramiding of *Rpv1, Rpv3*.*1*, and *Rpv10* achieved field-level resistance comparable to total immunity, without detectable fitness costs under standard viticultural conditions. This finding reinforces pyramiding as an effective route to achieve high levels and stability of resistance (Mundt, 2018), and complement integrated pest management approaches to reduce fungicide reliance of viticulture (Merdinoglu et al., 2018; Trapp et al., 2025; Zhang et al., 2025). However, the durability of grapevine resistance depends not only on genetic pyramiding but also on deployment strategies in vineyards. Modeling studies have shown that mixing single-gene and pyramided cultivars in the same landscape can accelerate resistance breakdown (Zaffaroni et al., 2024). However, our field plots established since 2004, indicate that certain pyramided combinations can remain stable even after two decades of downy mildew exposure, despite the presence of monogenic genotypes in proximity. These results support the possible field resilience and evolutionary advantage of stacking multiple resistance genes to buffer against pathogen adaptation (McDonald & Linde, 2002; Mundt, 2018), although furthers investigations are needed to identify the most effective spatial deployment strategies of new varieties carrying pyramided *Rpv*. Anyway, genetic resistances represent a natural and fragile capital and only an integrated approach that combines resistance diversity, agronomic practices, crop scouting and plant protection products can ensure a sustainable pathogen control while avoiding the emergence of resistance-breaking pathogen strains (Possamai et al., 2026).

### Broader significance and perspectives

The 2024 epidemic provided a rare, real-world stress test for the INRAE-ResDur breeding program. The remarkable consistency of resistance ranking across genotypes and populations, despite diverse genetic backgrounds, ages, and vineyard environments, underscores the robustness and broad transferability of the ResDur pyramiding strategy. However, some genotypes carrying pyramided *Rpv* still exhibited high downy mildew susceptibility, highlighting the need of combining phenotyping and genotyping techniques to validate the level of resistance of new varieties.

Future INRAE breeding efforts will prioritize the integration of less used and unbroken *Rpv* loci as *Rpv2* (Marsan et al., 2025) and *Rpv8* (Blasi et al., 2011) with the already implemented ones to provide grape varieties with new and, likely more durable, *Rpv* pyramiding combinations.

In parallel, the progress in genomics opens opportunities to shed light on grapevine quantitative resistance (QR) components, like constitutive defenses (e.g., secondary plant metabolites), induced defense responses (e.g., cell wall thickening), pathogen/damage associated patterns (PAMP/DAMP)-triggered resistance, enhanced signal transduction, and loss of genes required for susceptibility (Nelson et al., 2018; Pilet-Nayel et al., 2017; Poland et al., 2009). Disentangling the polygenic architecture of QR through genomic selection, GWAS, and multi-locus prediction models would enable breeders to combine major gene with alternatives, durables, broad-spectrum resistances, thereby reducing the risk of resistance breakdown and enhancing long-term field resilience (Nelson et al., 2018). This integrative approach, merging major genes with genomic prediction of QR, represents a promising new direction for conventional resistance breeding in grapevine.

Advances in new genetic resistance discovery and introduction, marker-assisted selection, genomic prediction pipelines (Bettinelli et al., 2023) and resistance phenotyping (Possamai et al., 2025) are expected to accelerate the development of next-generation resistant cultivars, ensuring both field efficacy and durability under evolving pathogen pressures. The INRAE-ResDur breeding program thus provides a foundational model for implementing such strategies, bridging classical resistance gene pyramiding with modern genomic tools to achieve sustainable viticulture with minimal fungicide input.

## Conclusion

Under historically severe downy mildew pressure, pyramiding *Rpv1, Rpv3*.*1*, and *Rpv10* conferred highly effective and stable resistance, far exceeding the protection achieved by single loci. The field data clearly demonstrate additive interactions between loci and support pyramiding as the cornerstone strategy for durable grapevine resistance breeding. These results provide a practical benchmark for integrating genetic resistance into sustainable viticulture systems and validate the long-term vision of the INRAE-ResDur breeding program.

## Acknowledgements

The authors thank all collaborators who have contributed to the INRAE-ResDur breeding program since its inception in the early 2000s. We are also grateful to the technical staff of the Unité Expérimentale Agronomique et Viticole (UE0871, INRAE–Centre Grand Est–Colmar, France) for their dedicated management and maintenance of the experimental vineyard plots.

## Authors’ contributions

GA conceived and designed the study. GA and ES conducted the phenotyping, and GA performed the statistical analyses. EP managed the genotyping work. TP and KA supervised the overall research. GA, TP, SW and KA wrote the manuscript. All authors contributed to the manuscript’s revision and approved the final version.

## Data availability statement

All the data are included in the article and its supplementary files.

## Supplementary Files

### Supplementary File S1. Details of the SSR markers utilized in the *Rpv* genotyping

Resistance loci to *P. viticola* (*Rpv*) segregating in the INRAE-ResDur populations, genetic SSR markers utilized for the assisted selection, and alleles of the parental plants of the populations. Resistance associated alleles are bold or underlined.

### Supplementary File S2. Summary of plant material evaluated in the study

Number of genotypes representing each INRAE-ResDur populations (50001, 42050, 50035 and mixed genotypes) and *Rpv* combinations of the study.

### Supplementary File S3. Cultural units of INRAE-ResDur populations in Colmar (France)

### Supplementary File S4. Downy mildew damages on INRAE-ResDur populations

Number of genotypes and descriptive statistics of leaf area affected by downy mildew (%) for the different grapevine populations (50001, 42050, 50035 and mixed population) and *Rpv* combinations of the study.

### Supplementary FileS5. Downy mildew damages on grapevine foliage

Different percentages of leaf damaged by downy mildew (defoliation, dessication of the leaf blade, sporulation of the pathogen and necrosis associated with the grapevine defence response) according to the scores utilized to assess the grapevine genotypes of the study (one averages core for four vines).

### Supplementary File S6. Downy mildew damages on INRAE-ResDur *Rpv* combinations

Number of genotypes and descriptive statistics of leaf area affected by downy mildew (%) for the mains *Rpv* combinations of the INRAE-ResDur programme.

### Supplementary File 7

Dataset. Leaf damaged by downy mildew epidemic (%) for each grapevine genotype (average score for four vine) of the INRAE-ResDur populations assessed on the 2 September 2024.

## References

Avia, K., Schneider, C., Onimus, C., Arnold, G., Dumas, V., Umar-Faruk, A., Butterlin, G., Dorne, M.-A., Alais, A., Jaegli, N., Lacombe, M.-C., Piron, M.-C., Prado, E., Wiedemann-Merdinoglu, S., Mestre, P., Duchêne, É., & Merdinoglu, D. (2023, luglio 17). The French grapevine breeding program ResDur: State of the art and perspectives. 2nd GiESCO, Cornell University, Jul 2023, Ithaca (Cornell University), United States. https://hal.inrae.fr/hal-04231604

Barker, C. L., Donald, T., Pauquet, J., Ratnaparkhe, M. B., Bouquet, A., Adam-Blondon, A.-F., Thomas, M. R., & Dry, I. (2005). Genetic and physical mapping of the grapevine powdery mildew resistance gene, Run1, using a bacterial artificial chromosome library. Theoretical and Applied Genetics, 111(2), 370–377. 10.1007/s00122-005-2030-8

Bellin, D., Peressotti, E., Merdinoglu, D., Wiedemann-Merdinoglu, S., Adam-Blondon, A.-F., Cipriani, G., Morgante, M., Testolin, R., & Di Gaspero, G. (2009). Resistance to Plasmopara viticola in grapevine ‘Bianca’ is controlled by a major dominant gene causing localised necrosis at the infection site. Theoretical and Applied Genetics, 120(1), 163–176. 10.1007/s00122-009-1167-2

Bettinelli, P., Nicolini, D., Costantini, L., Stefanini, M., Hausmann, L., & Vezzulli, S. (2023). Towards Marker-Assisted Breeding for Black Rot Bunch Resistance: Identification of a Major QTL in the Grapevine Cultivar ‘Merzling’. International Journal of Molecular Sciences, 24(4), 3568. 10.3390/ijms24043568

Blasi, P., Blanc, S., Wiedemann-Merdinoglu, S., Prado, E., Rühl, E. H., Mestre, P., & Merdinoglu, D. (2011). Construction of a reference linkage map of Vitis amurensis and genetic mapping of Rpv8, a locus conferring resistance to grapevine downy mildew. Theoretical and Applied Genetics, 123(1), 43–53. 10.1007/s00122-011-1565-0

Bois, B., Zito, S., & Calonnec, A. (2017). Climate vs grapevine pests and diseases worldwide: The first results of a global survey. OENO One, 51(2), 133. 10.20870/oeno-one.2016.0.0.1780

Calonnec, A., Wiedemann-Merdinoglu, S., Delière, L., Cartolaro, P., Schneider, C., & Delmotte, F. (2013). The reliability of leaf bioassays for predicting disease resistance on fruit: A case study on grapevine resistance to downy and powdery mildew. Plant Pathology, 62(3), 533–544. 10.1111/j.1365-3059.2012.02667.x

Chambre d’agriculture d’Alsace et Fredon Grand Est. (2024). Bulletin de Santé du Végétal: Viticulture. N° 14 novembre 2024, pp. 2–10. https://grandest.chambres-agriculture.fr/fileadmin/user_upload/273_chambre_dagriculture_grand_est/Agritheque/Collection_de_BSV/BSV/BSV_Viticulture_Alsace/2024/BSVBilan_VITI_ALS_2024.pdf

Chedid, E., Rustenholz, C., Avia, K., Dumas, V., Merdinoglu, D., & Duchêne, É. (2023). Impact of the introgression of resistance loci on agro-œnological traits in grapevine interspecific hybrids.

Delannoy, D., Maury, O., & Décome, J. (2022). CLIMATIK: Système d’information pour les données du réseau agroclimatique INRAE [Dataset]. Recherche Data Gouv. 10.57745/AJNXEN

Di Gaspero, G., Copetti, D., Coleman, C., Castellarin, S. D., Eibach, R., Kozma, P., Lacombe, T., Gambetta, G., Zvyagin, A., Cindrić, P., Kovács, L., Morgante, M., & Testolin, R. (2012). Selective sweep at the Rpv3 locus during grapevine breeding for downy mildew resistance. Theoretical and Applied Genetics, 124(2), 277–286. 10.1007/s00122-011-1703-8

European Parliament. (2021). The future of crop protection in Europe. Publications Office. 10.2861/086545

Fermaud, M., Smits, N., Merot, A., Roudet, J., Thiéry, D., Wery, J., & Delbac, L. (2016). New multipest damage indicator to assess protection strategies in grapevine cropping systems: An indicator of multipest damage in grapevine. Australian Journal of Grape and Wine Research, 22(3), 450–461. 10.1111/ajgw.12238

Fouillet, E., Delière, L., Chartier, N., Munier-Jolain, N., Cortel, S., Rapidel, B., & Merot, A. (2022). Reducing pesticide use in vineyards. Evidence from the analysis of the French DEPHY network. European Journal of Agronomy, 136, 126503. 10.1016/j.eja.2022.126503

Gadoury, D. M., Cadle-Davidson, L., Wilcox, W. F., Dry, I. B., Seem, R. C., & Milgroom, M. G. (2012). Grapevine powdery mildew (Erysiphe necator): A fascinating system for the study of the biology, ecology and epidemiology of an obligate biotroph. Molecular Plant Pathology, 13(1), 1–16. 10.1111/j.1364-3703.2011.00728.x

Gessler, C., Pertot, I., & Perazzolli, M. (2011). Plasmopara viticola: A review of knowledge on downy mildew of grapevine and effective disease management. Phytopathologia Mediterranea, 50(1):3–44. 10.14601/Phytopathol_Mediterr-9360

Innerebner, G., Roschatt, C., & Schmid, A. (2020). Efficacy of fungicide treatments on grapevines using a fixed spraying system. Crop Protection, 138, 105324. 10.1016/j.cropro.2020.105324

Marsan, L., Prado, E., Wiedemann-Merdinoglu, S., Schmidlin, L., Blanc, S., Delame, M., Schnee, S., Arti, B., Barnabé, G., Velt, A., Dumas, V., Merdinoglu, D., Rustenholz, C., & Mestre, P. (2025). Rpv2 is part of a cluster of NLRs specific to Vitis rotundifolia and confers total resistance to grapevine downy mildew. Theoretical and Applied Genetics, 138(8), 177. 10.1007/s00122-025-04959-z

McDonald, B. A., & Linde, C. (2002). Pathogen population genetic, evolutionary potential and durable resistance. Annual Review of Phytopathology, 40(1), 349–379. 10.1146/annurev.phyto.40.120501.101443

Merdinoglu, D., Schneider, C., Prado, E., Wiedemann-Merdinoglu, S., & Mestre, P. (2018). Breeding for durable resistance to downy and powdery mildew in grapevine. OENO One, 52(3), 203–209. 10.20870/oeno-one.2018.52.3.2116

Merdinoglu, D., Wiedeman-Merdinoglu, S., Coste, P., Dumas, V., Haetty, S., Butterlin, G., & Greif, C. (2003). Genetic analysis of downy mildew resistance derived from muscadinia. 603. 10.17660/ActaHortic.2003.603.57

Mestre, P.-F., Merdinoglu, D., Merdinoglu-Wiedemann, S., Calonnec, A. A., Deliere, L., & Delmotte, F. F. (2013). Vers une gestion durable de la résistance de la vigne au mildiou.

Mora, O., Berne, J.-A., Drouet, J.-L., Le Mouël, C., & Meunier, C. (2023). Foresight: European Chemical Pesticide-Free Agriculture in 2050. 10.17180/CA9N-2P17

Mundt, C. C. (2018). Pyramiding for Resistance Durability: Theory and Practice. Phytopathology®, 108(7), 792–802. 10.1094/PHYTO-12-17-0426-RVW

Nelson, R., Wiesner-Hanks, T., Wisser, R., & Balint-Kurti, P. (2018). Navigating complexity to breed disease-resistant crops. Nature Reviews Genetics, 19(1), 21–33. 10.1038/nrg.2017.82

Ogle, D., Doll, J., Wheeler, A., & Dinno, A. (2023). FSA: Simple Fisheries Stock Assessment Methods (Versione R package version 0.9.5) [Software]. https://CRAN.R-project.org/package=FSA

Paineau, M., Mazet, I. D., Wiedemann-Merdinoglu, S., Fabre, F., & Delmotte, F. (2022). The Characterization of Pathotypes in Grapevine Downy Mildew Provides Insights into the Breakdown of Rpv3, Rpv10, and Rpv12 Factors in Grapevines. Phytopathology®, 112(11), 2329–2340. 10.1094/PHYTO-11-21-0458-R

Pauquet, J., Bouquet, A., This, P., & Adam-Blondon, A.-F. (2001). Establishment of a local map of AFLP markers around the powdery mildew resistance gene Run1 in grapevine and assessment of their usefulness for marker assisted selection: Theoretical and Applied Genetics, 103(8), 1201–1210. 10.1007/s001220100664

Pelissier, D. R., Delmotte, D. F., Delière, M. L., Martínez, D. J. R., Marolleau, L., Mazet, M. I., Couture, D. C., Wiedemann-Merdinoglu, S., Fabre, D. F., & Miclot, A.-S. (2025). Multiple geographic breakdown events of the Rpv1–Rpv3.1 pyramided resistance in grapevine by Plasmopara viticola.

Peressotti, E., Wiedemann-Merdinoglu, S., Delmotte, F., Bellin, D., Di Gaspero, G., Testolin, R., Merdinoglu, D., & Mestre, P. (2010). Breakdown of resistance to grapevine downy mildew upon limited deployment of a resistant variety. BMC Plant Biology, 10(1), 147. 10.1186/1471-2229-10-147

Pertot, I., Caffi, T., Rossi, V., Mugnai, L., Hoffmann, C., Grando, M. S., Gary, C., Lafond, D., Duso, C., Thiery, D., Mazzoni, V., & Anfora, G. (2017). A critical review of plant protection tools for reducing pesticide use on grapevine and new perspectives for the implementation of IPM in viticulture. Crop Protection, 97, 70–84. 10.1016/j.cropro.2016.11.025

Pilet-Nayel, M.-L., Moury, B., Caffier, V., Montarry, J., Kerlan, M.-C., Fournet, S., Durel, C.-E., & Delourme, R. (2017). Quantitative Resistance to Plant Pathogens in Pyramiding Strategies for Durable Crop Protection. Frontiers in Plant Science, 8, 1838. 10.3389/fpls.2017.01838

Poland, J. A., Balint-Kurti, P. J., Wisser, R. J., Pratt, R. C., & Nelson, R. J. (2009). Shades of gray: The world of quantitative disease resistance. Trends in Plant Science, 14(1), 21– 29. 10.1016/j.tplants.2008.10.006

Possamai, T., Baltenweck, R., Wiedeman-Merdinoglu, S., Lacombe, M.-C., Dorne, M.-A., Bareyre, M., Griem, E., Fuchs, R., Bogs, J., Duchêne, E., Mestre, P., & Hugueney, P. (2025). Metabolic biomarker-based phenotyping unveils quantitative effects of plant resistance and pathogen aggressiveness in the grapevine (Vitis spp.)—Downy mildew (Plasmopara viticola) pathosystem. bioRxiv 2025.09.25.678557. 10.1101/2025.09.25.678557

Possamai, T., & Wiedemann-Merdinoglu, S. (2022). Phenotyping for QTL identification: A case study of resistance to Plasmopara viticola and Erysiphe necator in grapevine. Frontiers in Plant Science, 13, 930954. 10.3389/fpls.2022.930954

Possamai, T., Wiedemann-Merdinoglu, S., Lacombe, M.-C., Dorne, M.-A., Griem, E., Fuchs, R., Bogs, J., Baltenweck, R., Avia, K., Merdinoglu, D., & Hugueney, P. (2026). Defeated major resistance loci may act as Trojan horses compromising resistance pyramiding in grapevine. iScience, 114800. 10.1016/j.isci.2026.114800

R Core Team. (2023). R: A language and environment for statistical computing [Software]. .Http://www.R-project.org/

Rex, F., Fechter, I., Hausmann, L., & Töpfer, R. (2014). QTL mapping of black rot (Guignardia bidwellii) resistance in the grapevine rootstock ‘Börner’ (V. riparia Gm183 × V. cinerea Arnold). Theoretical and Applied Genetics, 127(7), 1667–1677. 10.1007/s00122-014-2329-4

Rockel. (2025). VIVC. Vitis International Variety Catalogue (May 2025) [Dataset]. https://www.vivc.de

Schneider, C., Onimus, C., Prado, E., Dumas, V., Wiedemann-Merdinoglu, S., Dorne, M. A., Lacombe, M. C., Piron, M. C., Umar-Faruk, A., Duchêne, E., Mestre, P., & Merdinoglu, D. (2019). INRA-ResDur: The French grapevine breeding programme for durable resistance to downy and powdery mildew. Acta Horticulturae, (1248), 207–214. 10.17660/ActaHortic.2019.1248.30

Schwander, F., Eibach, R., Fechter, I., Hausmann, L., Zyprian, E., & Töpfer, R. (2012). Rpv10: A new locus from the Asian Vitis gene pool for pyramiding downy mildew resistance loci in grapevine. Theoretical and Applied Genetics, 124(1), 163–176. 10.1007/s00122-011-1695-4

Trapp, O., Avia, K., Borrelli, C., Eibach, R., Merdinoglu, D., & Töpfer, R. (2025). More sustainability in Europe’s vineyards – Using resistant grapevine varieties to reduce the input of pesticides. PLANTS, PEOPLE, PLANET, 7(6), 1621–1628. 10.1002/ppp3.70038

Vezzulli, S., Malacarne, G., Masuero, D., Vecchione, A., Dolzani, C., Goremykin, V., Mehari, Z. H., Banchi, E., Velasco, R., Stefanini, M., Vrhovsek, U., Zulini, L., Franceschi, P., & Moser, C. (2019). The Rpv3-3 Haplotype and Stilbenoid Induction Mediate Downy Mildew Resistance in a Grapevine Interspecific Population. Frontiers in Plant Science, 10, 234. 10.3389/fpls.2019.00234

Welter, L. J., Göktürk-Baydar, N., Akkurt, M., Maul, E., Eibach, R., Töpfer, R., & Zyprian, E. M. (2007). Genetic mapping and localization of quantitative trait loci affecting fungal disease resistance and leaf morphology in grapevine (Vitis vinifera L). Molecular Breeding, 20(4), 359–374. 10.1007/s11032-007-9097-7

Wickham, H. (2016). Data Analysis. In H. Wickham, Ggplot2 (pp. 189–201). Springer International Publishing. 10.1007/978-3-319-24277-4_9

Wiedemann-Merdinoglu, S., Lacombe, M. C., Dorne, M. A., Dumas, V., Onimus, C., Prado, E., Schneider, C., Louise Dit Adèle, S., Misbach, J., Negrel, L., Baltenweck, R., Hugueney, P., & Merdinoglu, D. (2022). Fine monitoring of the effects of grapevine resistance loci on the development of Plasmopara viticola. BIO Web of Conferences, 50, 02005. 10.1051/bioconf/20225002005

Zaffaroni, M., Papaïx, J., Geffersa, A. G., Rey, J.-F., Rimbaud, L., & Fabre, F. (2024). Combining Single-Gene-Resistant and Pyramided Cultivars of Perennial Crops in Agricultural Landscapes Compromises Pyramiding Benefits in Most Production Situations. Phytopathology®, 114(10), 2310–2321. 10.1094/PHYTO-02-24-0075-R

Zendler, D., Töpfer, R., & Zyprian, E. (2020). Confirmation and Fine Mapping of the Resistance Locus Ren9 from the Grapevine Cultivar ‘Regent’. Plants, 10(1), 24. 10.3390/plants10010024

Zhang, Z., Niu, Z., Chen, Z., Zhao, Y., & Yang, L. (2025). Review of the Pathogenic Mechanism of Grape Downy Mildew (Plasmopara viticola) and Strategies for Its Control. Microorganisms, 13(6), 1279. 10.3390/microorganisms13061279

