## Supplementary Files_1 to 6 for "Pyramided resistance ensures grapevine (Vitis spp.) protection during high downy mildew (Plasmopara viticola) epidemic pressure"

| *Rpv* factor | SSR marker | 3082-1-42 | | Regent | | 3160-11-3 | | Bronner | | Artaban | | Divico | |
|---|---|---|---|---|---|---|---|---|---|---|---|---|---|
| | | allele 1 | allele 2 | allele 1 | allele 2 | allele 1 | allele 2 | allele 1 | allele 2 | allele 1 | allele 2 | allele 1 | allele 2 |
| *Rpv1* | VMC4f3.1 | 187 | **189** | 187 | 189 | 178 | **191** | 172 | 207 | **189** | 189 | 207 | 207 |
| *Rpv1* | VMC8g9 | **156** | 160 | 171 | 174 | **156** | 174 | 171 | 174 | **156** | 171 | 174 | 174 |
| *Rpv3.1*/ <u>*Rpv3.3*</u> | SC8_0096_022 | 201 | 215 | 201 | **220** | 204 | 215 | 202 | <u>222</u> | 201 | **220** | 217 | <u>222</u> |
| *Rpv3.1*/ <u>*Rpv3.3*</u> | UDV737 | 279 | 294 | **279** | 296 | 294 | 296 | <u>271</u> | 286 | **279** | 279 | <u>271</u> | 294 |
| *Rpv3.1*/ <u>*Rpv3.3*</u> | VMC7f2 | 195 | 197 | 201 | **207** | 197 | 199 | <u>197</u> | 197 | 195 | **207** | <u>197</u> | 197 |
| *Rpv10* | GF09-44 | 242 | 243 | 236 | 243 | 236 | 236 | **230** | 244 | 242 | 243 | **230** | 245 |
| *Rpv10* | GF09-47 | 293 | 293 | 287 | 293 | 293 | 293 | 293 | **296** | 287 | 293 | 293 | **296** |

**Supplementary File S1. Details of the SSR markers utilized in the *Rpv* genotyping.** Resistance loci to P. viticola (Rpv) segregating in the INRAE ResDur populations, genetic SSR markers utilized for the assisted selection, and alleles of the parental plants of the populations. Resistance associated alleles are bold or underlined.

| ***Rpv* combination** | **ResDur1 50001** | **ResDur2 42050** | **ResDur3 50035** | **Mixed population** | **Genotypes per *Rpv* combination** |
|---|---|---|---|---|---|
| *Rpv-* | 16 | 0 | 0 | 0 | 16 |
| *Rpv1* | 9 | 8 | 0 | 0 | 17 |
| *Rpv3.1* | 11 | 0 | 0 | 0 | 11 |
| *Rpv1/3.1* | 14 | 0 | 0 | 9 | 23 |
| *Rpv1/3.3* | 0 | 19 | 11 | 0 | 30 |
| *Rpv1/10* | 0 | 27 | 11 | 0 | 38 |
| *Rpv1/3.1/3.3* | 0 | 0 | 0 | 15 | 15 |
| *Rpv1/3.1/10* | 0 | 0 | 11 | 17 | 28 |
| *Rpv1/3.3/10* | 0 | 41 | 13 | 18 | 72 |
| *Rpv1/3.1/3.3/10* | 0 | 0 | 18 | 18 | 36 |
| **Genotypes per population** | 50 | 95 | 64 | 77 | **Total** 286 |

**Supplementary File S2. Summary of plant material evaluated in the study.** Number of genotypes representing each INRAE ResDur populations (50001, 42050, 50035 and mixed genotypes) and Rpv combinations of the study.

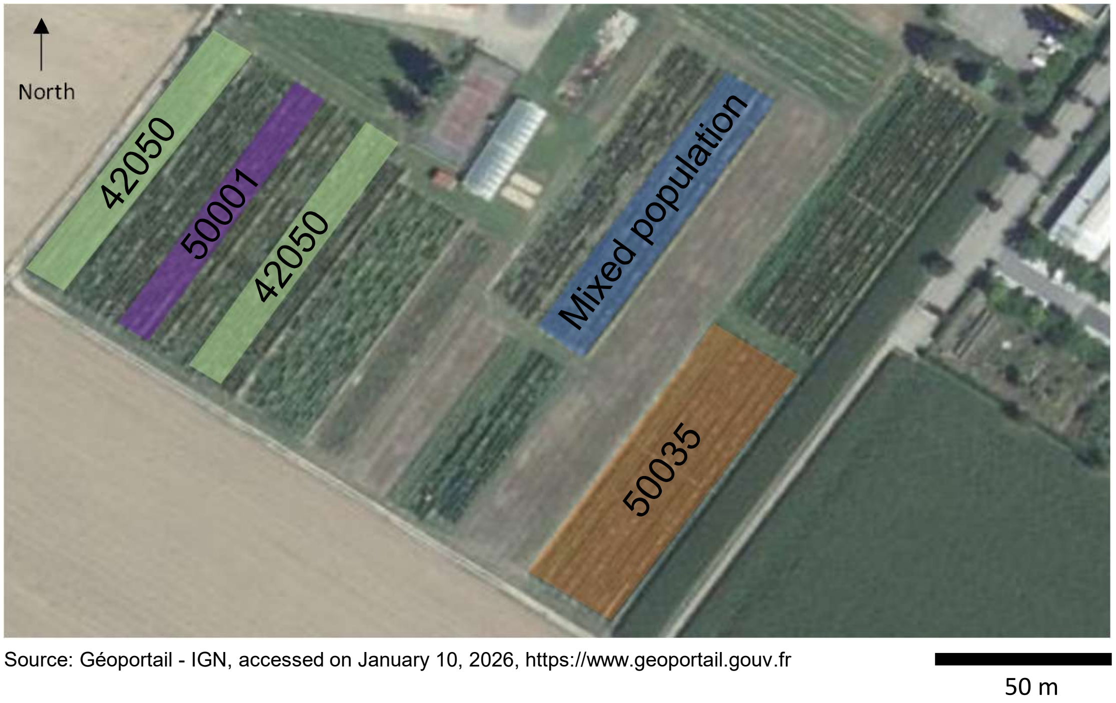


Source: Géoportail - IGN, accessed on January 10, 2026, https://www.geoportail.gouv.fr

**Supplementary File S3. Cultural units of INRAE ResDur populations in Colmar (France).**

| Population | *Rpv* combination | N° of genotypes | Leaf damaged by downy mildew | |
|---|---|---|---|---|
| | | | Mean% | Standard deviation% |
| ResDur1 50001 | *Rpv-* | 16 | 66.6 | 14.7 |
| | *Rpv3.1* | 11 | 48.2 | 12.0 |
| | *Rpv1* | 9 | 24.4 | 10.4 |
| | *Rpv1/3.1* | 14 | 20.4 | 12.2 |
| ResDur2 42050 | *Rpv1* | 8 | 45.0 | 19.2 |
| | *Rpv1/3.3* | 19 | 30.9 | 22.4 |
| | *Rpv1/3.3/10* | 41 | 11.6 | 15.7 |
| | *Rpv1/10* | 27 | 7.9 | 12.1 |
| ResDur3 50035 | *Rpv1/3.3* | 11 | 28.0 | 19.9 |
| | *Rpv1/10* | 11 | 16.3 | 16.8 |
| | *Rpv1/3.3/10* | 13 | 11.1 | 12.4 |
| | *Rpv1/3.1/3.3/10* | 18 | 3.1 | 7.2 |
| | *Rpv1/3.1/10* | 11 | 2.3 | 2.9 |
| Mixed population | *Rpv1/3.3/10* | 18 | 18.3 | 15.4 |
| | *Rpv1/3.1* | 9 | 18.4 | 13.5 |
| | *Rpv1/3.1/3.3* | 15 | 10.0 | 16.4 |
| | *Rpv1/3.1/3.3/10* | 18 | 6.3 | 8.6 |
| | *Rpv1/3.1/10* | 17 | 5.6 | 9.0 |

**Supplementary File S4. Downy mildew damages on INRAE ResDur populations.** Number of genotypes and descriptive statistics of leaf area affected by downy mildew (%) for the different grapevine populations (50001, 42050, 50035 and mixed population) and Rpv combinations of the study.

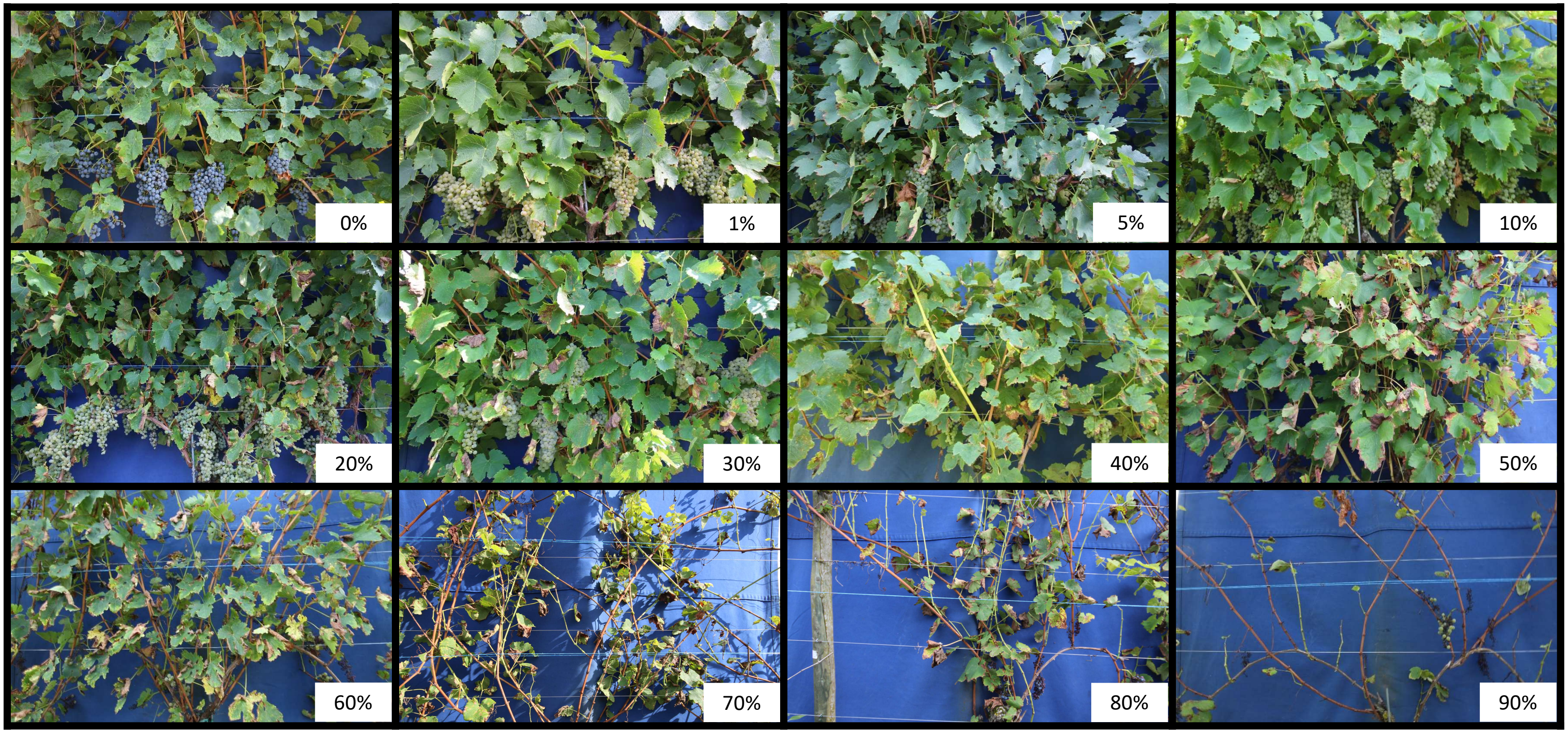


**Supplementary File S5.** Different percentages of leaf damaged by downy mildew (defoliation, dessication of the leaf blade, sporulation of the pathogen and necrosis associated with the grapevine defence response) according to the scores utilized to assess the grapevine genotypes of the study (one average core for four vines).

| *Rpv* combination | N° of genotypes | Leaf damaged by downy mildew | |
|---|---|---|---|
| | | Mean% | Standard deviation% |
| *Rpv-* | 16 | 66.6 | 14.7 |
| *Rpv3.1* | 11 | 48. 2 | 12.7 |
| *Rpv1* | 47 | 31.5 | 20.2 |
| *Rpv1/3.1* **ResDur1** | 38 | 16.1 | 14.6 |
| *Rpv1/10* **ResDur2** | 110 | 12.3 | 14.8 |
| *Rpv1/3.1/10* **ResDur3** | 64 | 4.9 | 7.7 |

**Supplementary File S6. Downy mildew damages on INRAE ResDur *Rpv* combinations (excluding Rpv3.3).** Number of genotypes and descriptive statistics of leaf area affected by downy mildew (%) for the mains Rpv combinations of the INRAE-ResDur programme.
